# Extended numerical analysis of an eyeball injury under direct impact

**DOI:** 10.1101/2021.02.26.433021

**Authors:** M. Koberda, A. Skorek, P. Kłosowski, M. Żmuda-Trzebiatowski, K. Żerdzicki, P. Lemski, U. Stodolska-Koberda

## Abstract

The objective of this study was to develop a numerical model of the eyeball and orbit to simulate a blunt injury to the eyeball leading to its rupture, as well as to conduct a comparative analysis of the results obtained using the finite element method against the clinical material concerning patients who had suffered an eyeball rupture due to a blunt force trauma. Using available sclera biometric and strength data, a numerical model of the eyeball, the orbital contents, and the bony walls were developed from the ground up. Then, eight different blunt force injury scenarios were simulated. The results of numerical analyses made it possible to identify possible locations and configurations of scleral rupture. The obtained results were compared against the clinical picture of patients hospitalized at the Clinic of Ophthalmology in 2010–2016 due to isolated blunt force trauma to the eyeball. It has been demonstrated that the extent of damage observed on the numerical model that indicated a possible location of eyeball rupture did not differ from the clinically observed configurations of the scleral injuries, however this applies to injuries the extend of which did not exceed 2–3 clock-hours on the eyeball. It has been found that the direction of the impact applied determines the location of eyeball rupture. Most often the rupture occurs at the point opposite to the clock-hour of the impact application. The eyeball rupture occurs in the first 7–8 milliseconds after the contact with the striking object, assuming that it strikes at a speed close to the speed of a human fist. It has been established that the injuries most often affected the upper sectors of the eyeball. Men are definitely more likely to sustain such injuries. Eyeball ruptures lead to significant impairment of visual acuity being most often degraded to light perception in front of the eye. Despite immediate specialist treatment, it is possible to achieve, on average, a visual acuity of 0.26 as per the Snellen scale. This study may contribute to a better understanding of injury mechanisms and better treatment planning. It may also contribute to the development of eyeball protection methods for persons exposed to the above-mentioned injuries.

## 1. Introduction

Each case of vision impairment resulting from injury has serious consequences both in terms of the quality of life of the injured person and from the socio-economic point of view. According to the World Health Organization (WHO), around 55 million eye injuries occur worldwide each year, of which about 1.6 million (about 3%) cases result in blindness (Négrel and Thylefors, 1998). According to these statistics, the largest percentage of injuries is blunt force trauma, followed by injuries from a sharp object, traffic accidents, gun shooting, injuries involving pyrotechnics, and falls (MacEwen et al., 1999; May et al., 2000). The worst prognosis is for injuries resulting from assault/beating and injuries sustained in motorcycle accidents (Liggett et al., 1990). Additionally, it has been demonstrated that the state under the influence of alcohol is not only conducive to the occurrence of serious eye injury but also tends to worsen the prognosis when the injury occurs (Jian-Wei et al., 2015). Injuries in children are a separate clinical problem. Among them, a particularly important mechanism that disrupts the scleral continuity is a penetrating trauma with a sharp object as well as blunt force injuries. The most common cases included accidental mutilation, violence, and communication injuries (Lesniak et al., 2012). Many of such injuries could have been avoided with proper eye protection.

Smith and Regan proposed two theories to describe the pathomechanism of orbital injury in the course of impact with a blunt object, i.e. the buckling theory and the hydraulic theory (Smith and Regan, 1957). The buckling theory assumes an orbital wall fracture when the force is applied to the outer rim. On the other hand, the hydraulic theory explains the formation of fractures of the lower and/or medial orbital wall, while the orbital wall remains intact. According to this theory, forces acting on the orbital soft tissues move them backward, leading to the creation of pressure which is transferred evenly to all orbital walls, and their rupture usually occurs in the thinnest place (Fujino and Makino, 1980; Skorek et al., 2014). The study by Ahmad et al. proves that such an injury mechanism protects the eyeball from rupture as a result of blunt trauma (Ahmad et al., 2003). In that study, it was found that less force was needed to break the orbital floor when the force was applied directly to the eyeball than when it was applied to the orbital rim. The orbital contents together with the eyeball and bony walls are closely interrelated in terms of mechanics. Most of the studies on eyeball resistance to injuries concerned models of the eyeball alone (Cui et al., 2015; Gray et al., 2011; Rossi et al., 2011; Stitzel et al., 2002). Accurate mapping of the orbital contents influences the value of results of modeling dynamic blunt trauma both to the eyeball and the orbit, as well as the conclusions drawn from such experiments (Kłosowski et al., 2019).

## 2. Objective of the study

Studies on the mechanisms of blunt trauma, its course, and its effect bring us closer to the determination of more effective methods of eye protection. For obvious reasons, such studies cannot take place in vivo, which is why they are increasingly often conducted on numerical models. This study aimed to present some aspects of numerical modeling of the eyeball and orbit and the elements of analysis of simulated blunt injury caused by a moving solid object. The results obtained with simulation methods were compared against clinical studies of patients with eyeball rupture hospitalized at the Department of Ophthalmology at the Medical University of Gdańsk.

## 3. Material and method

For the needs of the analysis, the properties of individual parts of the orbit and eyeball (except for the sclera) were adopted as per literature (Table 1). For the sclera, the elastic modulus (Young’s modulus) value of 10.7 MPa was adopted, being the average from the previously conducted own tests (on an animal model) (Kłosowski et al., 2019). Then, the orbital numerical model was constructed. The possibility of contact between the following elements was determined: the modeled solid body and the outside of the eyeball, the eyeball and the orbital fat, and the fat and the bone. Also, a membrane representing the orbital septum was created. From a mechanical point of view, it was assumed that both the eyeball interior the orbital fat were incompressible materials, which reflects the Poisson modulus value close to 0.5, and that bones, eyeball, and adipose tissue were elastic isotropic materials. Adoption of the Poisson modulus value *v* = 0.49 for incompressible elements caused a divergence of calculations. Only the value of *v* = 0.499999 leads to correct results (Żerdzicki et al., 2019; Zmuda Trzebiatowski and Skorek, 2020).

**Table 1:** Material properties adopted for the individual elements of the eyeball and orbit model (Dai et al., 2016)(1); (Uchio et al., 1999)(2); (Skorek et al., 2014)(3); (Kłosowski et al., 2019) (4))

| | Young's modulus $E$ [Pa] | Poisson modulus $\nu$ | Density $\rho$ [kg/m<sup>3</sup>] |
| --- | --- | --- | --- |
| Orbital bones (3) | $1,3 \times 10^9$ | 0.33 | 2000 |
| Vitreous body (1,2) | $0,5 \times 10^6$ | 0,499999 | 1000 |
| Orbital fat (4) | $0,5 \times 10^6$ | 0,499999 | 952 |
| Sclera and cornea (1,4) | $10 \times 10^6$ | 0,42 | 1000 |
| Orbital septum (4) | $0,5 \times 10^6$ | 0,33 | 1000 |

To obtain the most reliable and life-like results, in the numerical analysis a geometrically nonlinear variant in the range of large deformations was used in the Lagrange approach. In the Msc.Marc / Mentat program (MSC Software, Newport Beach, California, USA), dynamic calculations were performed in the dynamic implicit options. The equations of motion were integrated using the Houbolt method with the parameters *γ*_1_ = 1.5 and *γ* = −0.5. The integration step was Δ*t* = 10 ^-6^ s. Moreover, it was assumed that all materials are isotropic and have elastic (bones) or elastic perfect plastic (soft tissues) physical properties (Table 1). Nevertheless, the analysis was terminated when the elastic limit of the sclera was reached at a certain point. It was assumed that it is the eyeball rupture point.

The obtained results were displacements, stress components, and equivalent stresses Huber-von Mises-Hencky σM calculated according to the formula:

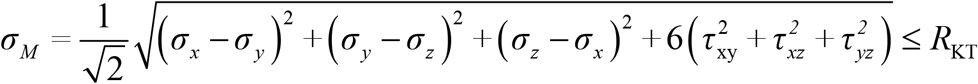

where: *σ*_*x*_, *σ*_*y*_, *σ*_*z*_ − normal stress consistent with the axes of the global coordinate system; *τ*_*xy*_, *τ*_*xz*_, *τ*_*yz*_ - shear stress in the same system; *R*_*KT*_ - ultimate stress value causing sclera rupture

The application of this hypothesis is because only one parameter is known, determining the moment of sclera rupture. In the obtained analyses, no stress component exceeded the permissible value of R_KT_, but the already calculated equivalent stresses *σ*_*M*_ exceeded these permissible values, and it was possible to end the simulation at this stage and recognize that the sclera ruptured.

The sclera and cornea were modeled as separate parts of the eyeball since they feature different properties. For simplicity, eye lens was not included in the modeling due to their complicated suspension system and a minor role they could play in eyeball rupture.

The phenomenon of contact was taken into account in the calculations. For this purpose, four deformable bodies were defined: bone, eyeball, orbital septum and fat, and a single non-deformable body (or “punch”) that caused the system deformation. To build our model (Fig. 1), we partly adopted equations for curvatures of individual eyeball structures included in the work (Nogueira et al., 2011).

**Fig. 1:**
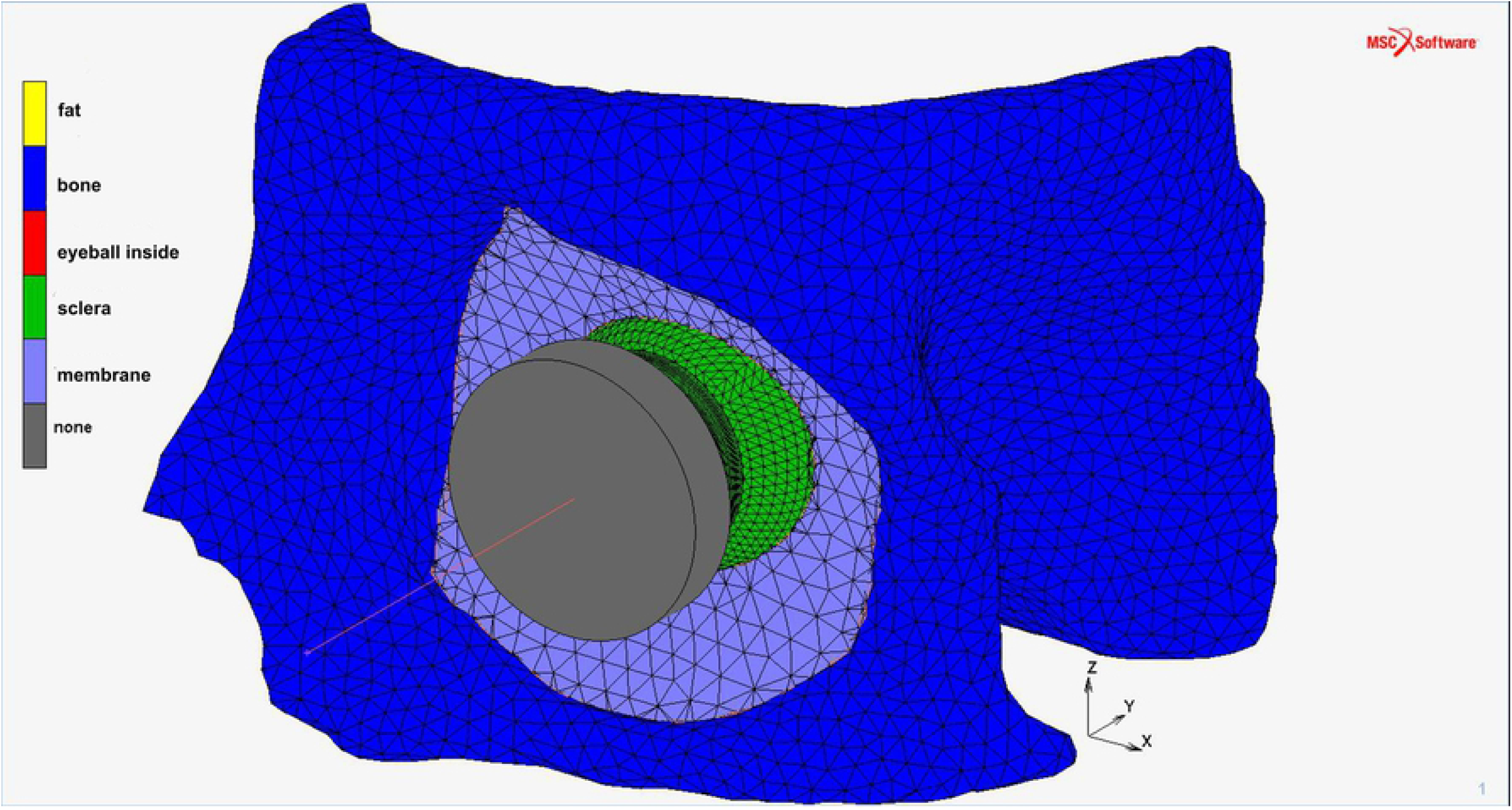
Finished rnodel of the eyeball, orbit and striking object.

First, the behavior of the numerical model of the eyeball was tested using a non-linear dynamic compression test on a rigid surface (Fig. 2).

**Fig. 2:**
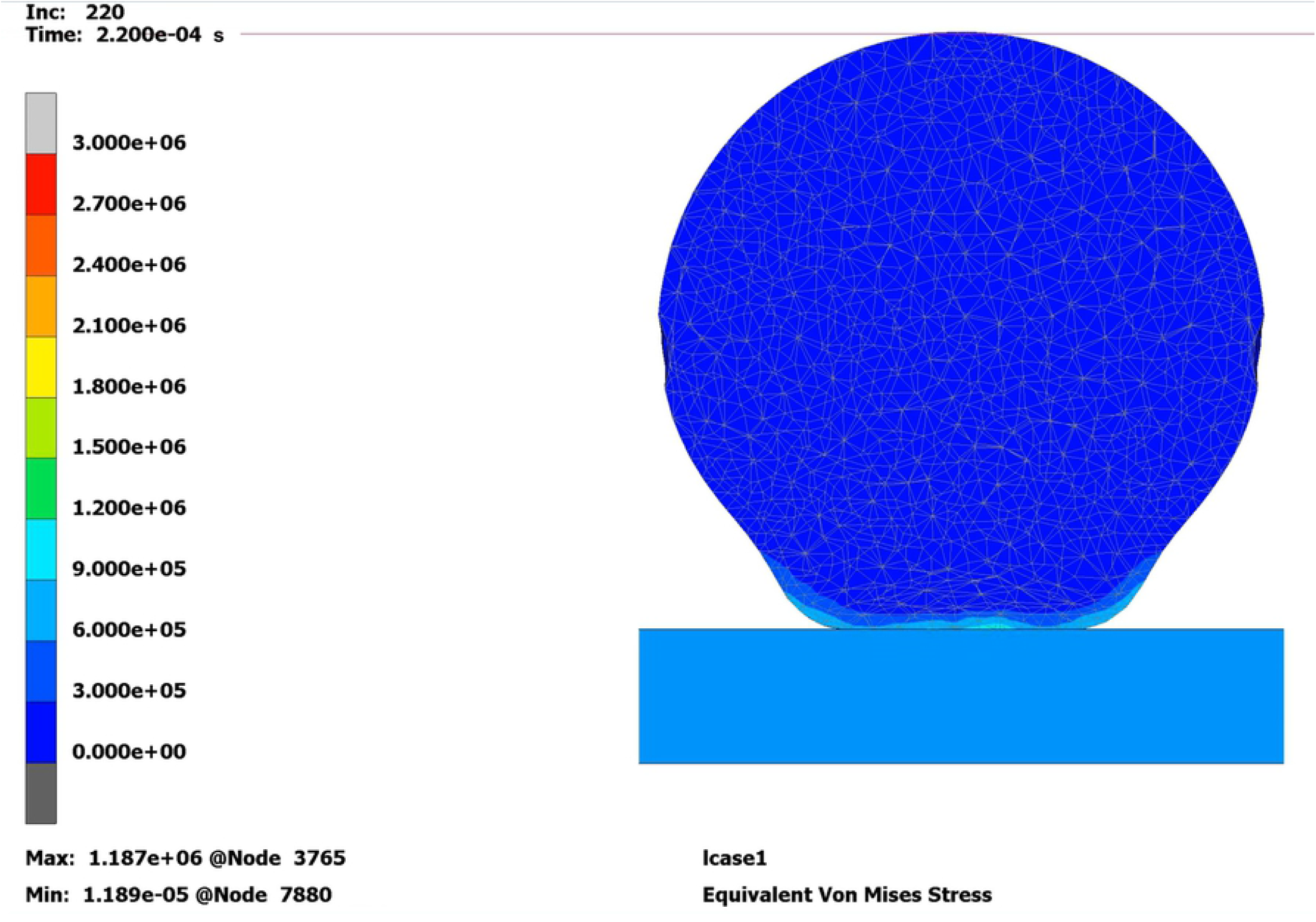
Test of the eyeball nurnerical model (Pa].

Our model consists of 22,673 nodes and 98,064 elements (of which 32,346 for the eyeball). The finished model of the eyeball was connected to the other elements of the orbit and a cylindrical solid object was modeled. This object was to strike at the eyeball directly at a speed of 9 m/s and then decelerate linearly to 0 m/s after 0.003 s. Eight tests were performed from eight different directions. Grid generation using the finite element method is shown in (Fig. 1). Velocity vectors for the specified impact directions are presented in Table 2.

**Table 2:**
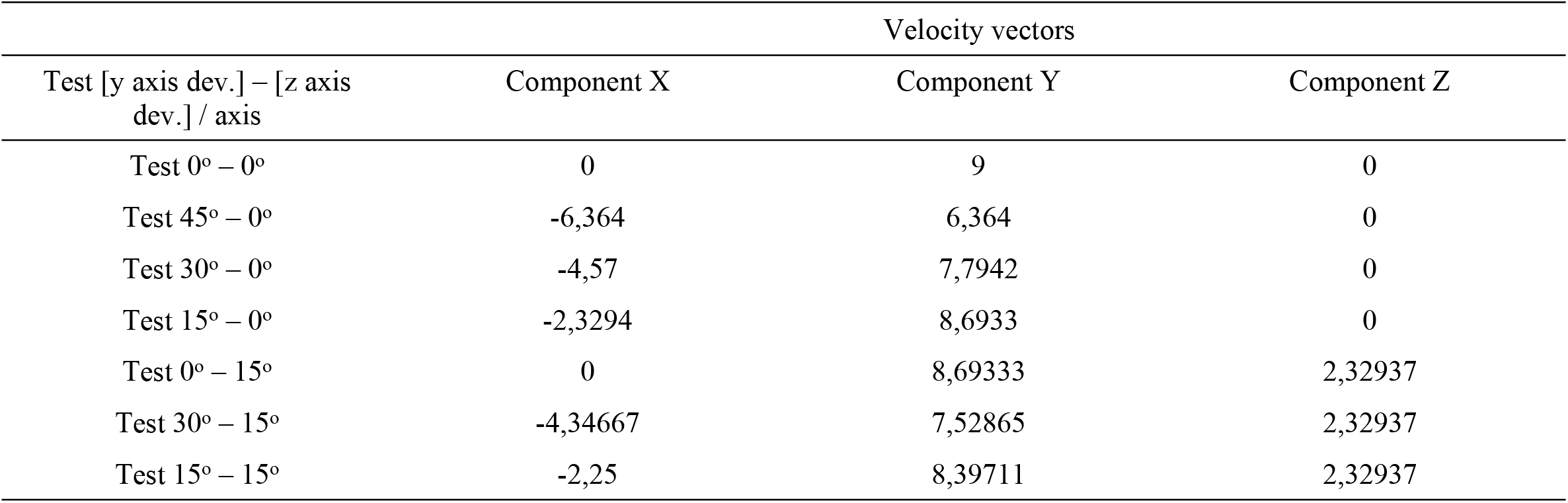
List of initial velocity vectors (in m/s) for the individual tests

| Test [y axis dev.] – [z axis dev.] / axis | Velocity vectors |  |  |
| --- | --- | --- | --- |
|  | Component X | Component Y | Component Z |
| Test 0° – 0° | 0 | 9 | 0 |
| Test 45° – 0° | -6,364 | 6,364 | 0 |
| Test 30° – 0° | -4,57 | 7,7942 | 0 |
| Test 15° – 0° | -2,3294 | 8,6933 | 0 |
| Test 0° – 15° | 0 | 8,69333 | 2,32937 |
| Test 30° – 15° | -4,34667 | 7,52865 | 2,32937 |
| Test 15° – 15° | -2,25 | 8,39711 | 2,32937 |

## 4. Results

### 4.1 FEM modeling of the eyeball rupture

The solution for finite element equations was obtained in the form of stress maps in the numerical model of the eyeball. It should be noted that all of the tests were performed for the left eyeball and left orbit model. In each test, the non-deformable body stroke at the eyeball from a different direction. The numerical model indicated the points where the assumed limit stress was achieved, identical to the most likely point of scleral rupture in vivo (Table 3). Following other authors’ reports, for the limit stresses resulting in scleral rupture the following values were adopted: [*strain*] ε = 6.8%; [*stress*] *R*_*K*_ = 9.4 MPa (Uchio et al., 1999). Similar values are found in (Stitzel et al., 2002).

**Table 3:** Summary of tests on the numerical model

| No. | Impact direction | Reference point where the limit stress was reached | Clock-hour where the limit stress was reached |
| --- | --- | --- | --- |
| Test No. 1<br>0° – 0° | From the front (along axis Y) | sclera in the place of the external rectus muscle attachment | 3:00 |
| Test No. 2<br>45° – 0° | From the front at an angle of 45° from axis Y (from the front and the temple) | sclera in the nasal quadrant in the projection of the flat part of the ciliary body (pars plana), slightly below the height of the internal rectus muscle attachment | 08:30 |
| Test No. 3<br>30° – 0° | From the front at an angle of 30° from axis Y (from the front and the temple) | sclera in the nasal quadrant in the projection of the flat part of the ciliary body (pars plana), slightly below the height of the internal rectus muscle attachment | 08:30 |
| Test No. 4<br>15° – 0° | From the front at an angle of 15° from axis Y (from the front and the temple) | sclera in the nasal quadrant in the projection of the flat part of the ciliary body (pars plana), slightly below the height of the internal rectus muscle attachment | 08:30 |
| Test No. 5<br>0° – 15° | From the front along axis Y and from below at an angle of 15° from axis Z (from below) | sclera in the upper quadrant, in the pars plana projection, in the projection of the upper rectus muscle attachment | 11:00 – 12:00 |
| Test 6<br>30° – 15° | From the front at an angle of 30° from axis Y and from below at an angle of 15° from axis Z (from the temple and from below) | sclera in the upper quadrant, in the pars plana projection, slightly nasally from the upper rectus muscle attachment | 11:00 – 12:00 |
| Test No. 7<br>15° – 15° | From the front at an angle of 15° from axis Y and from below at an angle of 15° from axis Z (from the temple and from below) | sclera in the upper quadrant, the pars plana projection, in two places: slightly nasally and slightly temporally from the projection of the upper rectus muscle attachment | 11:00 and 01:00 |

### 4.2 Comparison with clinical data

Then, the locations of eyeball rupture in patients hospitalized in the Ophthalmology Clinic of Medical University of Gdańsk in 2010–2016 were determined, as illustrated in Table 4.

**Table 4:** Summary of clinical cases

| No | Sex | Age | Eyeball | Mechanism of trauma | Rupture location | Rupture extent |
| --- | --- | --- | --- | --- | --- | --- |
| 1 | M | 67 | Left | Impact (wood fragment) | Corneal limbus | 6:00 to 10:00 |
| 2 | M | 78 | Left | Fall (on the pot) Impact (beating) | Muscle attachment + to the equator | 09:00 to 12:00 |
| 3 | M | 41 | Left | Impact (beating) | Corneal limbus | 04:00 |
| 4 | M | 59 | Right | Impact (beating) | Muscle attachment | 12:00 to 3:00 |
| 5 | M | 62 | Left | Fall (onto heater) | Corneal limbus | 10:00 to 12:00 |
| 6 | M | 61 | Right | Impact (wood trunk) | Corneal limbus + meridian wound | 11:00 to 6:00 |
| 7 | M | 67 | Right | Strike (sole) | Corneal limbus + oblique wound | 10:00 – 02:00 |
| 8 | F | 70 | Right | Fall (on bathroom faucet) | Corneal limbus | 11:00 to 4:00 |
| 9 | M | 58 | Right | Fall (getting off the bus) | Equator | 09:00 to 12:00 (90 degrees in the upper temporal quadrant) |
| 10 | M | 64 | Left | Bash (metal object) | Corneal limbus + 2 meridian wounds | 01:00 to 4:00 + meridional wounds at 01:00 and 4:00 almost to n. II |
| 11 | M | 60 | Right | Impact (beating) | Equator towards the back of the eyeball | 12:00 to 3:00 obliquely towards the rear pole |
| 12 | M | 54 | Left | Impact (beating) | Corneal limbus | 10:00 to 02:30 |
| 13 | M | 65 | Right | Fall | Corneal limbus | 6:00 to 12:00 |
| 14 | M | 26 | Right | Impact (hit by the car) | Pars plana | 01:00 to 02:30 |
| 15 | F | 69 | Right | Impact (with wood) | Corneal limbus | 01:00 to 06:00 |
| 16 | F | 53 | Left | Impact (chair rod) | Corneal limbus + meridian | 6:00 to 12:00 and then to the upper rectus muscle |
| 17 | M | 60 | Left | Fall (edge of well) | Corneal limbus | 12:00 to 05:00 |
| 18 | M | 60 | Right | Impact (probably against the steering wheel) | Equator | 09:00 to 01:00 |
| 19 | M | 41 | Right | Impact (with a piece of wood) | Meridian from the limbus to the equator | 07:00 |
| 20 | M | 59 | Right | Bang (champagne cork) | Corneal limbus | 09:00 |
| 21 | F | 60 | Right | Fall | Corneal limbus + meridian wound | 07:00 to 12:00 |
| 22 | M | 54 | Right | Impact (beating) | Equator | 09:30 to 12:00 |
| 23 | M | 10 | Right | Impact (shovel) | Corneal limbus + oblique wound | 07:30 to 12:00 |
| 24 | M | 64 | Left | Impact (against the corner of the shelf) | Upper rectus muscle attachment | 12:00 |

The data obtained in the analysis of the clinical material (Table 4) were compared against the data obtained in the dynamic analysis on the numerical model (Table 3), and such configurations of eyeball rupture in clinical patients were selected that correspond to the results of the numerical simulation regarding the location of the probable rupture, as presented in Table 5. The selected ones were presented in the form of legible figures (Fig. 3, 4, 5, 6, 7, 8).

**Table 5:** Comparison of the tests on the numerical model with the clinical cases

| No. | Clock-hour where the limit stress $R_K$ was reached in the numerical model | Number of the patient with a similar rupture |
| --- | --- | --- |
| Test No. 1<br>($0^\circ - 0^\circ$) | 03:00 | 3, 20 |
| Tests Nos. 2, 3, 4<br>($45^\circ - 0^\circ$)<br>($30^\circ - 0^\circ$)<br>($15^\circ - 0^\circ$) | 08:30 | 1 |
| Test No. 5<br>($0^\circ - 15^\circ$) | 11:00 to 12:00 | 11, 24 |
| Test 6<br>($30^\circ - 15^\circ$) | 11:00 to 12:00 | 2, 4, 5, 8, 14 |
| Test No. 7<br>($15^\circ - 15^\circ$) | 11:00 and 01:00 | 7, 12 |

**Fig. 3:**
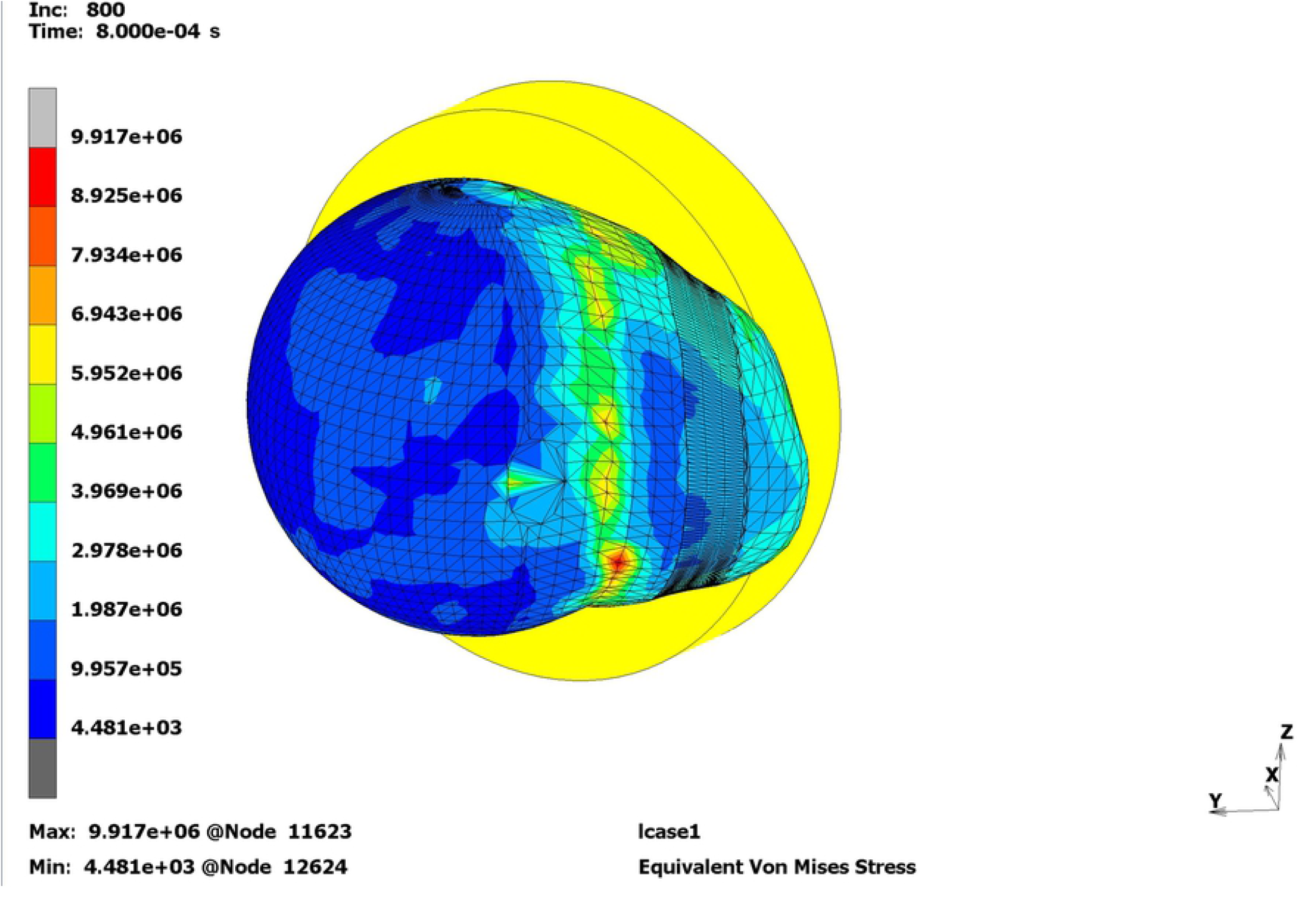
Predicted sci era rupture site in test No. 2 (Pa].

**Fig. 4:**
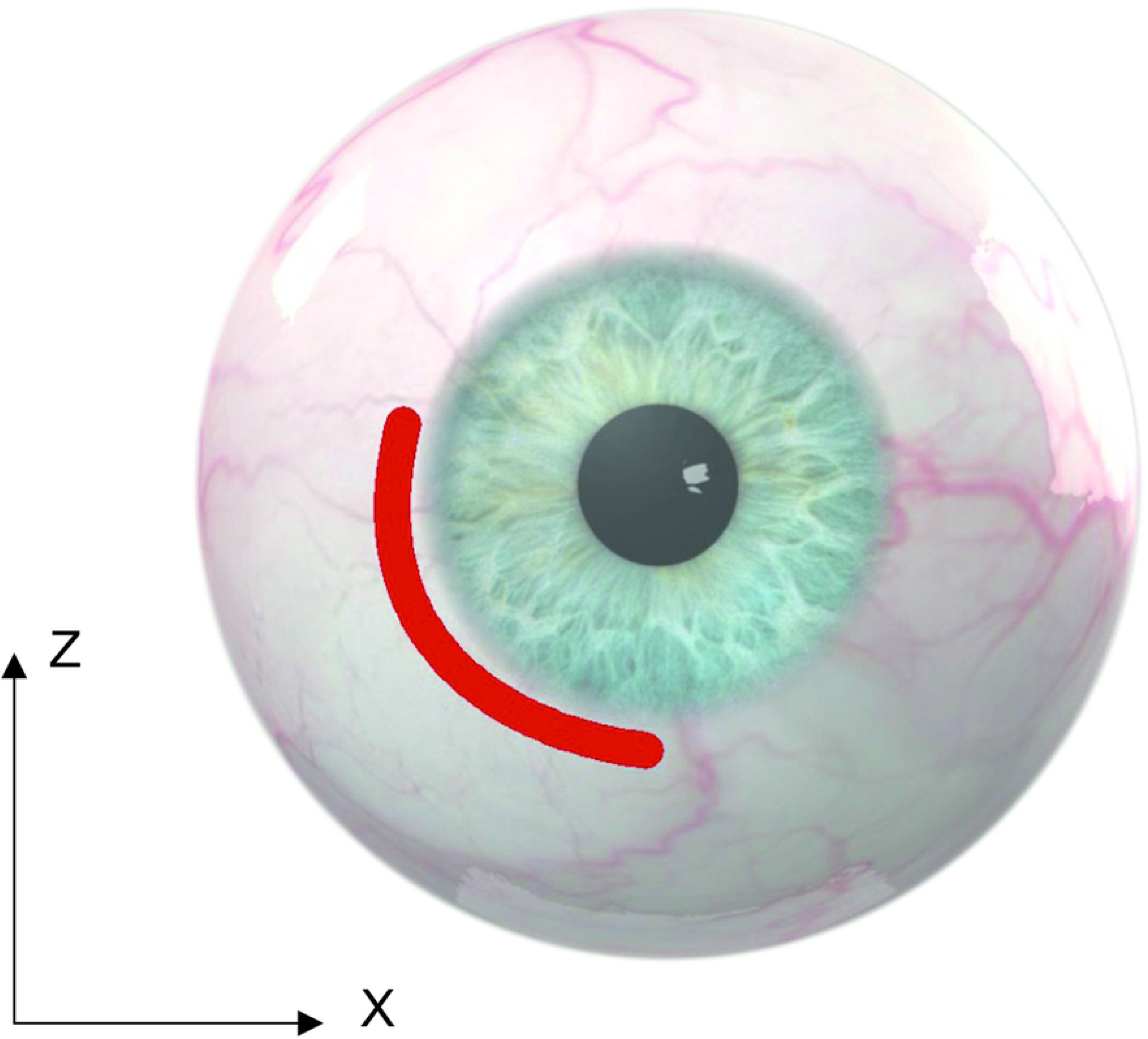
Location of the wound in patient No. I (left eye)

**Fig. 5:**
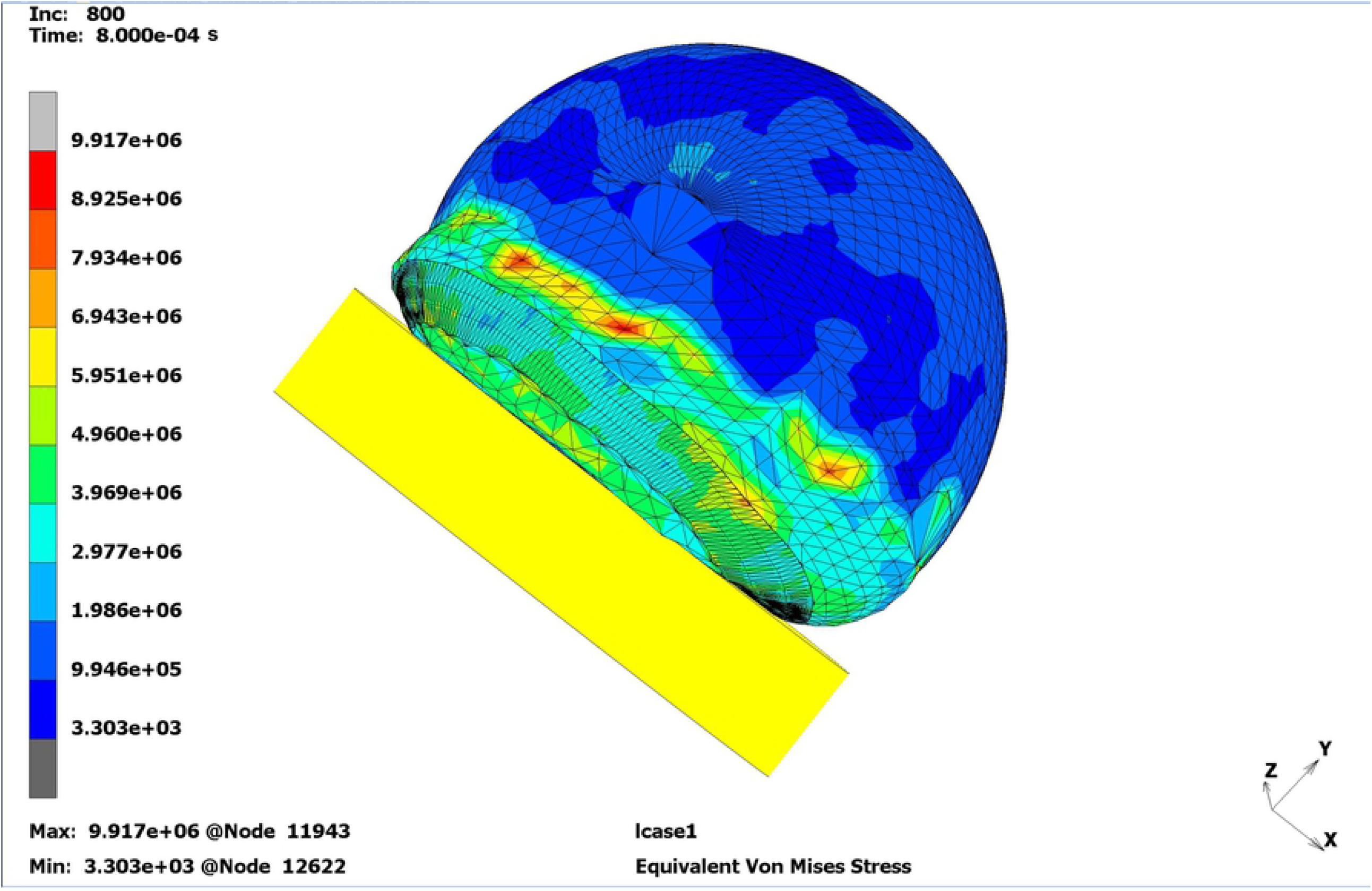
Predicted sclera rupture site in test No. 5 (Pa)

**Fig. 6:**
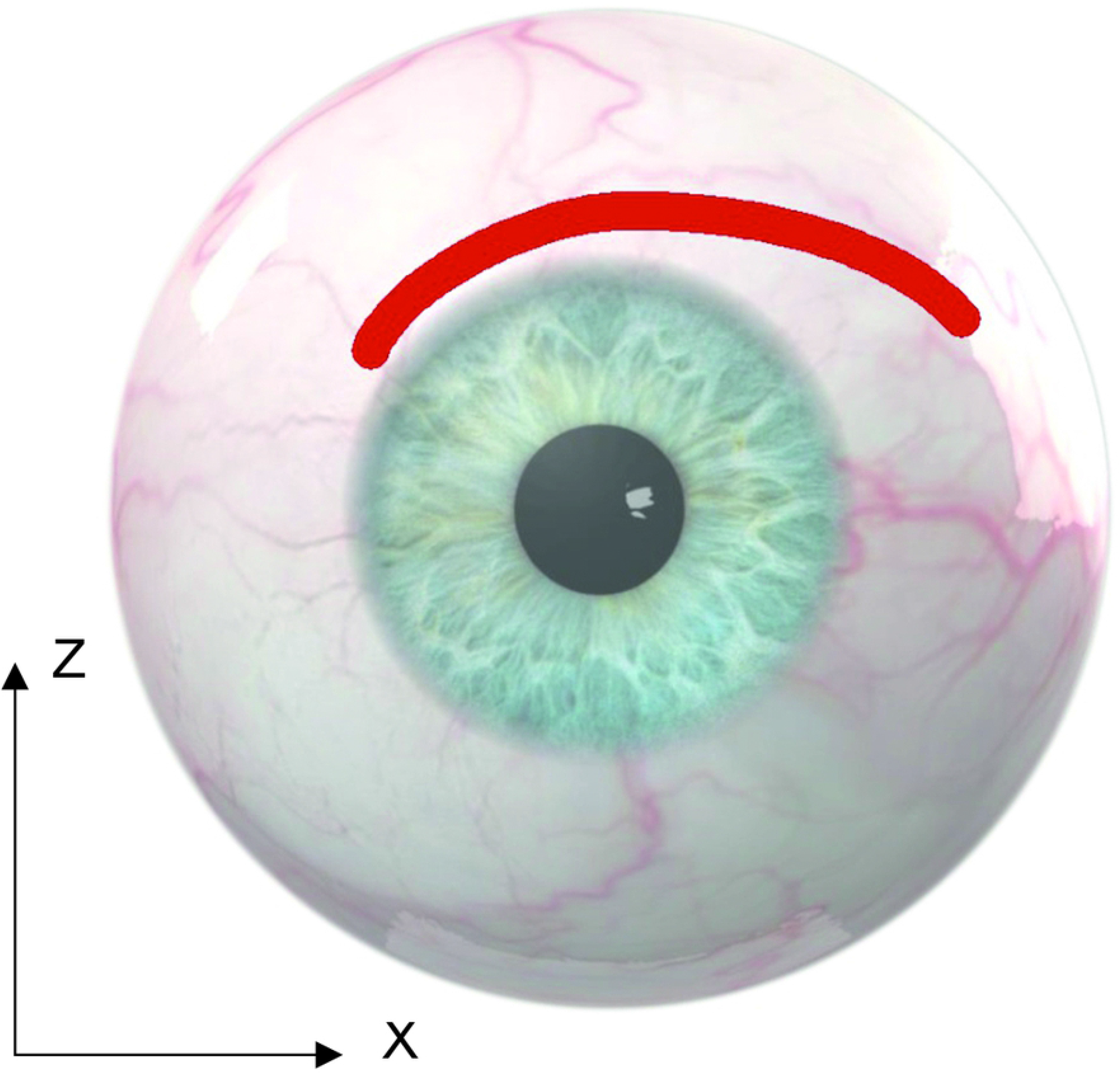
Location of the wound in patient No 7 (right eye)

**Fig. 7:**
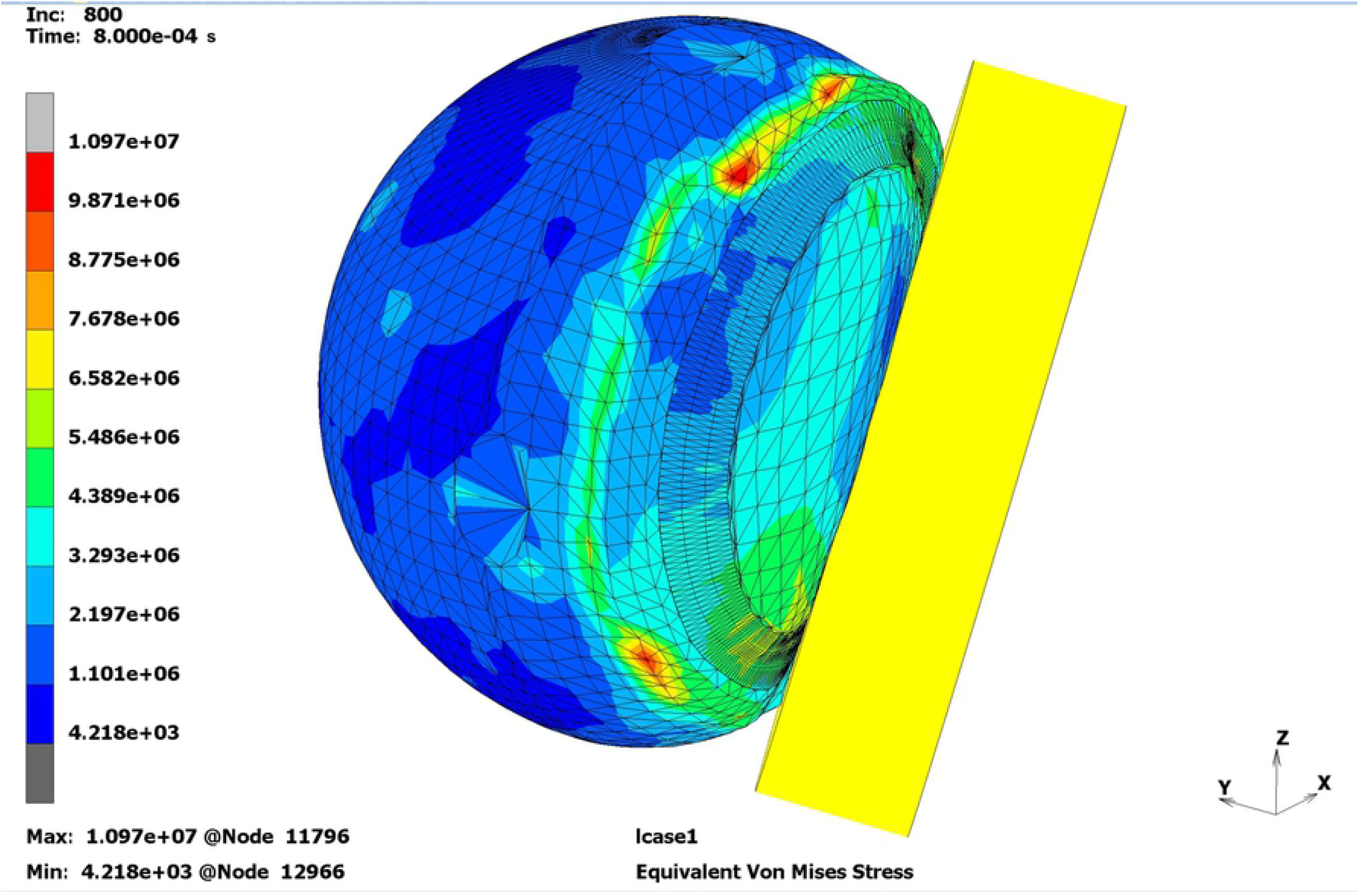
Predicted sclera rupture site in test No. 6 [Pa].

**Fig. 8:**
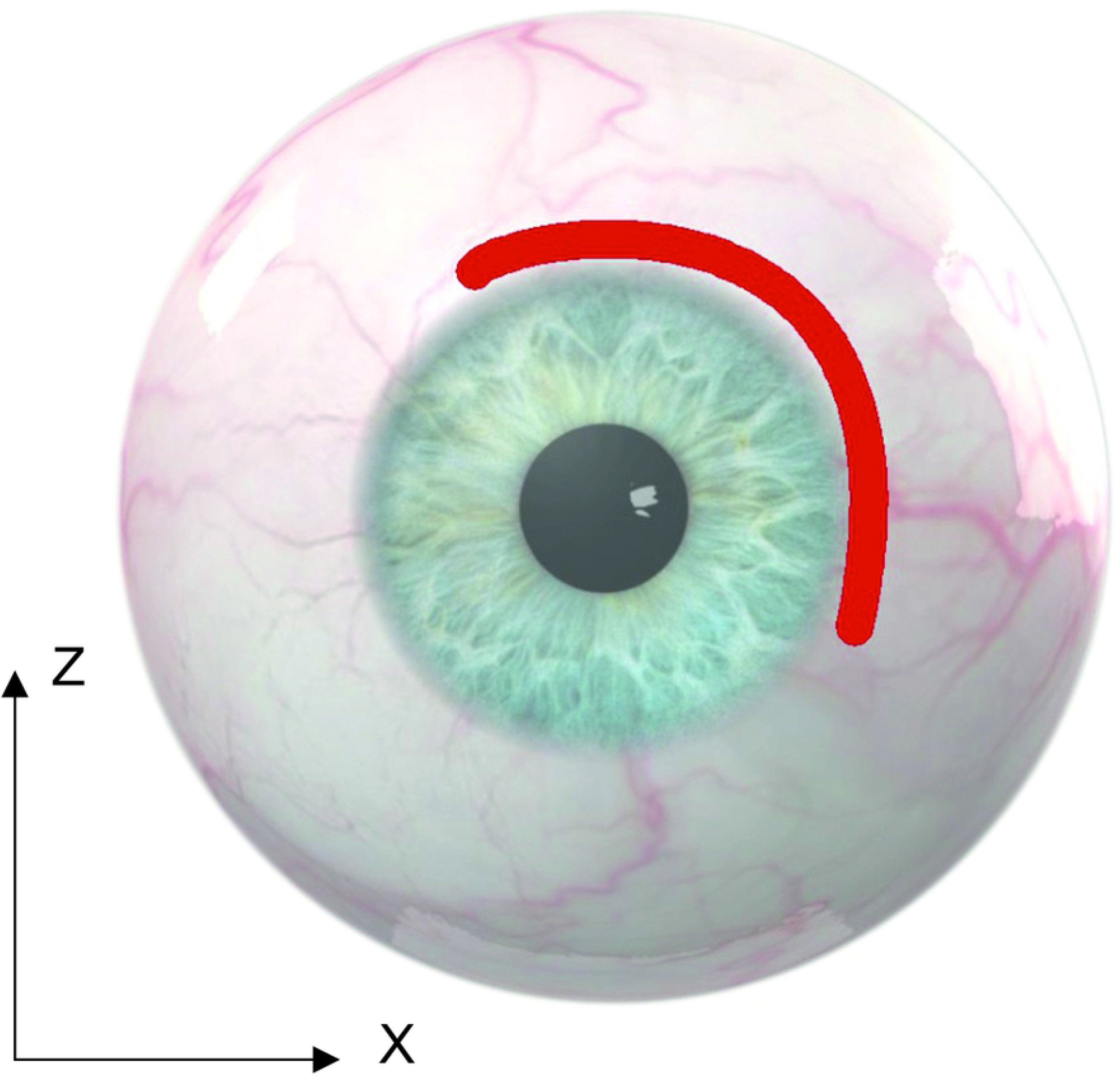
Location of the wound in patient No. 8 (right eye)

In half of the patients (No. 3, 6, 9, 10, 13, 15–19, and 21– 23 as per Table 4), no rupture configuration matching the locations indicated in the numerical simulation was observed, or the rupture size was too large to indicate the probable original rupture spot. Most likely, a mechanism of injury in these patients was complex, or the impact directions varied from those proposed in Table 3.

## 5. Discussion

The literature describes various objectives and applications for eyeball numerical models. Nagasao et al., Schaller et al., and Al-Shukhun et al. were the first ones to use mathematical methods (finite element methods) for orbital model construction. Initially, the models contained only bony elements. Later on, they were enriched with orbital contents (eyeball and other retrobulbar structures) (Al-sukhun et al., 2012; Al-Sukhun et al., 2011, 2006; Nagasao et al., 2010, 2009, 2006; Schaller et al., 2013). More advanced models were built based on the CT image transferred with the use of computational programs to a mathematical finite element model. The basis for the data transfer were grayscale (Hounsfield scale) units, considering only the linear coefficient for the weakening of density and not the material properties of the tissues. Those studies made it possible to obtain a new perspective on the traumatology of the orbit. For this reason, it was possible to assess the behavior of individual bone structures in the course of injury and find *locus minoris resistentie* of the orbit, as well as to determine particularly dangerous injury directions. From here, it is only a step to develop new protective measures for people who are particularly exposed to damage to the orbit, both in the workplace and during sports activities. However, the modeled orbital contents were treated as a kind of “filling” for bone structures, without high accuracy in the mapping of the eyeball, for example.

In the work by Gray et al., the objective was to compare the results of numerical modeling of an eye injury from a paintball impact with the empirical model of such an injury. In that work, an exact model of the eyeball was built, and consistency of the numerical model with the trauma course observed using high-speed cameras was demonstrated. Since ballistic tests require placing an examined eyeball in a block of transparent gel, the numerical model of the eye in that study was placed in similar conditions. Also, the authors demonstrated that numerical modeling of trauma could help in the understanding of the course and causes of intraocular injuries that have not yet been precisely explained (Gray et al., 2011). In the work by Han et al., the numerical model was used to further explain the mechanisms of drug transport from the eye surface to the anterior chamber, and the purpose of those studies was to contribute to improving the effectiveness of drugs administered in that way.

The innovation of our study lies in the fact that in none of the so-far published works a numerical model of the eyeball embedded in a numerical model of the orbit of a real patient was compared against clinical cases.

Studies on the mechanisms of blunt trauma, its course, and its effects certainly bring us closer to determining more effective methods of eye protection. For obvious reasons, such studies cannot take place in vivo, which is why they are increasingly often conducted using numerical models. In our work, some aspects were presented concerning numerical modeling of the eyeball and orbit as well as elements of the analysis of simulated blunt trauma caused by a moving solid object. The results obtained with simulation methods were compared against clinical studies of patients with eyeball rupture hospitalized at the Department of Ophthalmology at the Medical University of Gdańsk. The largest extent of damage was observed both in the numerical model and in patients in the cases of strokes presented as test No. 7 (15°/15°). It seems that this should be taken into account in studies on the construction of eye protection elements.

## 6. Conclusions

The tests performed suggest that in the case of a blunt impact directly at the eyeball with an object moving at a speed of about 9 m/s from a specific direction, the eyeball rupture will begin at a specific and repetitive location. This knowledge can be used in situations where the configuration of the eyeball rupture is known but it is necessary to determine the direction from which the impact might have occurred.

The results of the analysis on the created numerical model can be compared against real eyeball ruptures observed in patients in hospital conditions. In many cases, the rupture configuration is consistent with the simulated rupture sites; however, this comparison cannot be applied for extensive ruptures, but only for those limited to 2–3 clock-hours of the sclera. This is because for more extensive wounds it is not possible, at the current stage of research, to determine the direction of rupture widening, therefore it cannot be presumed which clock-hour of the rupture was the initial, and which one was the terminal point of the rupture.

The limit stress on the created numerical model was achieved no later than 8 ms from the contact of the object striking at the cornea of the modeled eyeball. This observation suggests that the sclera begins to rupture before the impact pulse stops acting.

Further development of the numerical model details to include periocular muscles and other so-far omitted structures, while considering the plasticity effect or, finally, including the damage phenomenon in simulations, will result in obtaining more accurate results and lead to drawing more far-reaching conclusions.

## 9. Method of Literature Search

The objective of our search strategy was to find past or recent articles that mention properties of porcine and/or human sclera, preparation of numerical model of the eye and/orbit, and blunt trauma of eye and orbit. For the literature review, we used standard search strategies involving the querying of online databases (mostly Pubmed) using keywords, followed by evaluation of the bibliographies of relevant articles.

